# Modeling the emergence of viral resistance for SARS-CoV-2 during treatment with an anti-spike monoclonal antibody

**DOI:** 10.1101/2023.09.14.557679

**Authors:** Tin Phan, Carolin Zitzmann, Kara W. Chew, Davey M. Smith, Eric S. Daar, David A. Wohl, Joseph J. Eron, Judith S. Currier, Michael D. Hughes, Manish C. Choudhary, Rinki Deo, Jonathan Z. Li, Ruy M. Ribeiro, Ruian Ke, Alan S. Perelson, the ACTIV-2/A5401 Study Team

**Affiliations:** Theoretical Biology & Biophysics, Los Alamos National Laboratory, Los Alamos, NM, USA; Department of Medicine, David Geffen School of Medicine, University of California, Los Angeles, CA, USA; Department of Medicine, University of California, San Diego, CA, USA; Lundquist Institute at Harbor-UCLA Medical Center, Torrance, CA, USA; Department of Medicine, University of North Carolina at Chapel Hill School of Medicine, Chapel Hill, NC, USA; Harvard T.H. Chan School of Public Health, Boston, MA, USA; Department of Medicine, Division of Infectious Diseases, Brigham and Women’s Hospital, Harvard Medical School, Boston, MA, USA; Santa Fe Institute, Santa Fe, NM, USA

**Keywords:** SARS-CoV-2, monoclonal antibody treatment, Bamlanivimab, viral rebound, viral dynamics

## Abstract

The COVID-19 pandemic has led to over 760 million cases and 6.9 million deaths worldwide. To mitigate the loss of lives, emergency use authorization was given to several anti-SARS-CoV-2 monoclonal antibody (mAb) therapies for the treatment of mild-to-moderate COVID-19 in patients with a high risk of progressing to severe disease. Monoclonal antibodies used to treat SARS-CoV-2 target the spike protein of the virus and block its ability to enter and infect target cells. Monoclonal antibody therapy can thus accelerate the decline in viral load and lower hospitalization rates among high-risk patients with susceptible variants. However, viral resistance has been observed, in some cases leading to a transient viral rebound that can be as large as 3-4 orders of magnitude. As mAbs represent a proven treatment choice for SARS-CoV-2 and other viral infections, evaluation of treatment-emergent mAb resistance can help uncover underlying pathobiology of SARS-CoV-2 infection and may also help in the development of the next generation of mAb therapies. Although resistance can be expected, the large rebounds observed are much more difficult to explain. We hypothesize replenishment of target cells is necessary to generate the high transient viral rebound. Thus, we formulated two models with different mechanisms for target cell replenishment (homeostatic proliferation and return from an innate immune response anti-viral state) and fit them to data from persons with SARS-CoV-2 treated with a mAb. We showed that both models can explain the emergence of resistant virus associated with high transient viral rebounds. We found that variations in the target cell supply rate and adaptive immunity parameters have a strong impact on the magnitude or observability of the viral rebound associated with the emergence of resistant virus. Both variations in target cell supply rate and adaptive immunity parameters may explain why only some individuals develop observable transient resistant viral rebound. Our study highlights the conditions that can lead to resistance and subsequent viral rebound in mAb treatments during acute infection.

**Author summary:** Monoclonal antibodies have been used as a treatment for SARS-CoV-2. However, viral evolution and development of variants has compromised the use of all currently authorized monoclonal antibodies for SARS-CoV-2. In some individuals treated with one such monoclonal antibody, bamlanivimab, transient nasal viral rebounds of 3-4 logs associated with resistant viral strains occur. To better understand the mechanisms underlying resistance emergence with high viral load rebounds, we developed two different models that incorporate drug sensitive and drug resistant virus as well as target cell replenishment and fit them to data. The models accurately capture the observed viral dynamics as well as the proportion of resistant virus for each studied individual with little variation in model parameters. In the models with best-fit parameters, bamlanivimab selects for resistance mutants that can expand to high levels due to target cell replenishment. The ultimate clearance of virus however depends on the development of adaptive immunity.

## Introduction

As of September 2023, SARS-CoV-2 has caused over 6.9 million deaths worldwide (WHO, 2022). While vaccines are available to reduce disease severity, a large fraction of the world’s population is not vaccinated (WHO, 2022a). With new variants that escape immunity rapidly emerging, effective treatment for severe cases is urgently needed (CDC, 2022a). Anti-SARS-CoV-2 monoclonal antibodies (mAbs) represent one class of effective treatments (Taylor et al., 2021). For example, the first monoclonal antibody that received emergency use authorization (EUA) for the treatment of SARS-CoV-2, bamlanivimab (BAM), can accelerate the viral decline and lower hospitalization and death rates among high-risk patients (Chew et al., 2022; Chen et al., 2021). One concerning observation during treatment of SARS-CoV-2 infection with mAbs is the emergence of viral resistance and concomitant viral rebound. In the case of BAM treatment, after the initial peak in the viral load, viral rebounds of up to 3-4 orders of magnitude consisting mostly of resistant virus have been observed in small subsets of patients in multiple studies (Choudhary et al., 2022; Jensen et al., 2021; Peiffer-Smadja et al., 2021). During rebound, infectious viruses are culturable, implying virus may be able to be transmitted and infect others (Boucau et al., 2022a).

Since mAbs are a treatment for SARS-CoV-2 and other viral infections such as respiratory syncytial virus and Ebola virus (Panteleo et al., 2022), understanding the mechanisms that allow resistant virus to emerge and rebound is important for designing effective treatment strategies that reduce disease severity and the chance of on-treatment viral transmission. Furthermore, such analyses can provide insights on the underlying dynamics of SARS-CoV-2 infection, such as the impact of the host target cell regeneration and adaptive immune responses on viral rebound, which has potential implications for the development of the next generation of mAb therapies and viral rebound after nirmatrelvir-ritonavir or other antiviral therapies (Boucau et al., 2022b; Carlin et al., 2022; Charness et al., 2022; Dai et al., 2022; Ranganath et al., 2022).

Here, we use data from the ACTIV-2/A5401 randomized controlled clinical trial (NCT04518410) of BAM (Chew et al., 2022; Choudhary et al., 2022) to investigate the mechanism that gives rise to the observed transient viral rebound associated with the emergence of resistant virus. BAM is an IgG1 monoclonal antibody that binds to the SARS-CoV-2 virus’s spike protein, blocking its ability to enter and infect target cells. Due to its high potency for neutralizing SARS-CoV-2 (Chew et al., 2022; Chen et al., 2021), BAM can select for escape mutations such as E484K, an emergent mutation observed in the majority of rebound cases (Choudhary et al., 2022; Jensen et al., 2021; Peiffer-Smadja et al., 2021). As a single substitution mutation, E484K is likely to be present at low frequencies at the time of treatment initiation and selected for in the presence of BAM (see text S1, Supplementary Material), yet only a small fraction of treated patients shows signs of resistance emergence and transient viral rebound (Choudhary et al., 2022). The selective pressure of BAM is likely the reason why some resistant mutations become dominant during the transient viral rebound; however, there must be other important factors associated with high viral load rebound.

Drug resistance to antiviral treatments has been studied extensively for many viral pathogens (Stilianakis et al., 1998; Kepler and Perelson, 1998; Ribeiro and Bonhoeffer, 2000; Rong et al., 2010; Ke et al., 2018). Rapid viral growth under treatment requires high resistance of the virus and the availability of abundant target cells for the resistant virus to replicate and spread further as discussed in Ke et al. (2018). Without enough target cells, the selected resistant virus is unlikely to generate an observable transient viral rebound. For SARS-CoV-2, it has been shown that the viral load peaks around the time of or a few days after symptom onset (Ke et al., 2021; Cevik et al., 2021; Killingley et al., 2022), suggesting cells that are available for early infection either become depleted (by infection) or protected by innate immune responses (Ke et al., 2021) causing the virus to fall post-peak. The fact that the emergence of resistant virus associated with high viral load rebound occurs several days post symptom onset in some patients treated with BAM (Choudhary et al., 2022) is, therefore, difficult to explain.

Mathematical models are well suited to study the phenomenon of drug resistance and viral rebound (Ribeiro and Bonhoeffer, 2000; Rong et al., 2010; Zhang et al., 2014; Ke et al., 2018; Perelson and Ke, 2021; Cardozo-Ojeda et al., 2021b; Perelson et al., 2023). Thus, to better understand the crucial factors leading to the emergence of resistant virus associated with high viral load rebound, we built models that include feasible mechanisms that can increase the availability of potential virus producing cells: the natural proliferation of target cells such as epithelial cells expressing the ACE-2 receptor (Fang et al, 2020) and the loss of protection from infection by the innate immune response (Voigt et al., 2016) and fit these models to data from individuals who experienced viral rebound due to the emergence of resistant virus during BAM treatment.

## Methods

### Data

We analyzed data from the cohort of individuals with SARS-CoV-2 infection and symptoms consistent with COVID-19 who were given a single infusion of 700 mg BAM in a phase 2 randomized, placebo-controlled trial (ACTIV-2/A5401 study) described in detail previously (Chew et al., 2022; Choudhary et al., 2022). Viral load and mutant frequency data were obtained from anterior nasal swabs collected by the participants daily from the time of the BAM infusion to day 14 and at days 21 and 28. There were eight individuals who showed viral rebound with emerging resistance mutations in this treatment group. Patient B2-10, who showed signs of transient viral rebound with mutation S494P, only had four data points above the quantifiable limit and only one data point associated with the S494P resistance. Hence, we focus our study on the other seven participants with more frequent data (B2-2 to B2-8), who exhibited transient viral rebound associated with resistant mutants.

### Models

Classical target cell limited models with a single viral population have been used to study within-host dynamics of SARS-CoV-2 (Wang et al., 2020; Hernandez-Vargas et al., 2020; Gonçalves et al., 2020; Perelson and Ke, 2021; Maisonnasse et al., 2021; Kim et al., 2021; Heitzman-Breen and Ciupe, 2022; Esmaeili-Wellman et al., 2023) and other diseases such as HIV and HCV (Perelson et al., 1996; Perelson, 2002). Since our data set contains data on both the viral load and the frequency of resistant mutants, we used a two-strain SARS-CoV-2 model to utilize both types of data (Rong et al., 2010). In this model, target cells (*T*) represent the epithelial cells expressing ACE-2 receptors that are susceptible to infection. The target cell dynamics is governed by

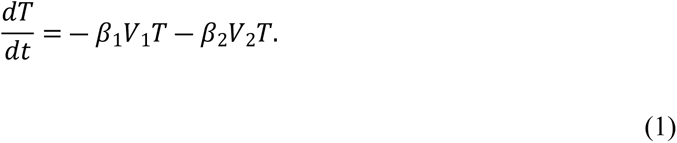

We classify SARS-CoV-2 viruses as either sensitive (*V*_1_) or resistant (*V*_2_) to BAM. Note that *V*_1_ and *V*_2_ do not indicate two particular genotypes of SARS-CoV-2, but rather groups of viruses that are either sensitive or resistant to BAM. The other equations defining the model are:

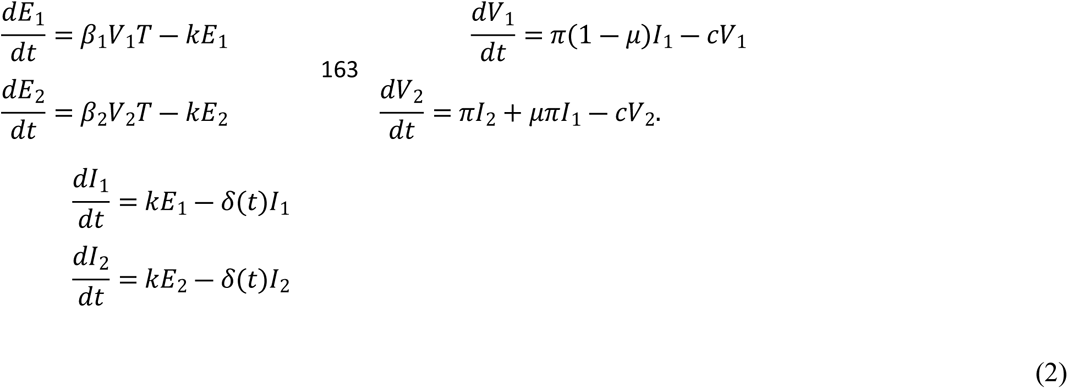

In this model, viruses *V*_1_ and *V*_2_ infect target cells at rate *β*_1_ and *β*_2_, respectively. The infection produces *E*_1_ and *E*_2_, which are cells infected by *V*_1_ and *V*_2_, respectively, in the eclipse phase of the viral lifecycle and not yet producing virus. After staying in the eclipse phase for an average duration of 1/*k*, infected cells start producing viruses at rate π and are reclassified as productively infected cells, *I*_1_ and *I*_2_, respectively. We also allow for a fixed mutation rate *μ* from the BAM sensitive to BAM resistant virus. Infected cells and viruses are cleared at per capita rates *δ*(*t*) and *c*, respectively. The loss rate of infected cells also includes the emergence of adaptive immunity, which is modeled as in the study by Pawelek et al. (2012) by

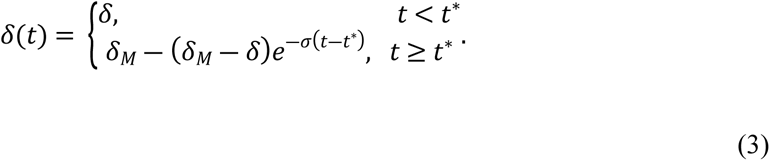

We define *δ* and *δ*_M_ as the baseline and maximum death rates of the infected cells, respectively. For simplicity, we set *δ_M_* = 10*δ*. Time *t*\* is when adaptive immunity starts to emerge and σ determines the emergent speed of adaptive immunity. Note that adaptive immunity affects the cells infected by either viral strain.

### Neutralizing effect and dynamics of neutralizing antibodies

We explicitly incorporate the passive infusion and decay dynamics of BAM based on the model developed by Cardozo-Ojeda and Perelson (2021). We let A(*t*) be the serum concentration of BAM with unit *μg mL*^−1^. We define *t*_inf_ as the time after infection that BAM infusion starts, Δ*T* as the duration of the infusion (assumed to be 1 hour or 1/24 day), and *A_max_* as the maximum serum concentration (taken from data for these seven study participants). The equation governing the antibody dynamics takes the form

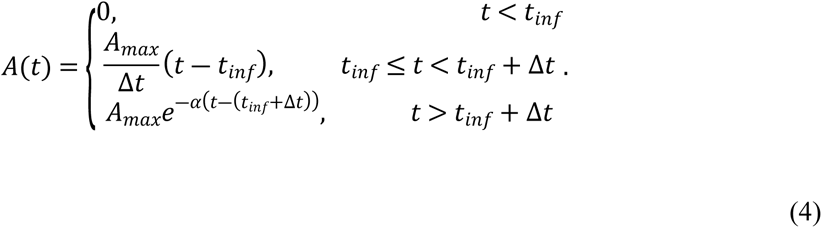

For each study participant, we have the maximum antibody concentration (*A_max_*) and the concentration at day 28 (*A*_28_). By fitting an exponential decay function that goes through the data points *A_max_* and *A*_28_ we estimate the decay rate *α* (Table S1, Supplementary Material). Note that we do not include a loss term for *A*(*t*) due to the binding to the viruses or a gain term when the antibody dissociates from an immune complex. This is a simplification based on the observation that 99% of individuals infused with BAM have mAb concentrations greater than the IC90 (the concentration required for 90% inhibition) 28 days after the infusion (Chew et al., 2022). Thus, antibodies are in great excess, and we assume the mAb concentration is not noticeably affected by interaction with the virus.

The neutralizing effect of BAM is modeled as follows. Antibodies bind to sensitive and resistant viruses at rates *k_on_*_,1_ and *k_on_*_,2_, respectively, to create neutralized virus-antibody immune complexes *C*_1_ and *C*_2_, respectively, which are no longer able to infect target cells. The complexes dissociate at rates *k_off_*_,1_ and *k_off_*_,2_ and are cleared at rates *γ*_1_ and *γ*_2_, respectively for the sensitive and resistant viruses. This leads to the following modifications to the basic target cell limited model.

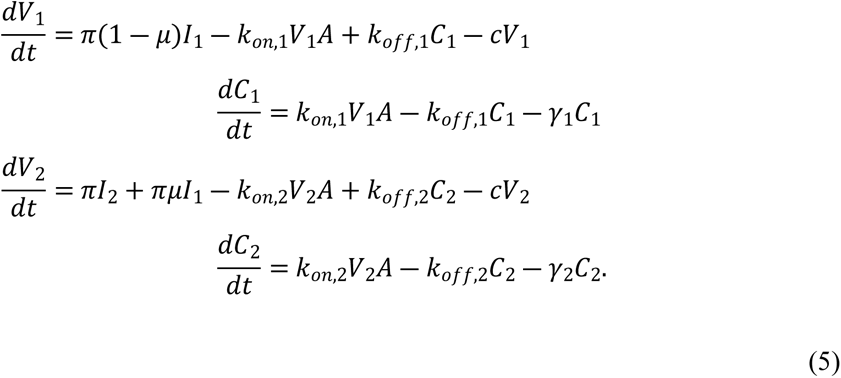

Together, equations (1) – (5) constitute the basic viral dynamic model with the neutralizing effect of antibodies. The E484K mutation, which is observed in 5 out of the 7 participants with transient viral rebound, is an escape mutant that abrogates BAM binding. A previous in vivo study showed that the binding affinity of BAM to the E484K spike protein is less than 1% of the binding to wild type SARS-CoV-2 (Chen et al., 2021). Hence, we assume *k_on_*_,2_ is negligible, and thus we do not account for the dynamics of *C*_2_.

Our first intuition is perhaps to attribute the observed viral rebound solely to resistance to BAM as the majority of rebound viruses are resistant mutants. However, the timing of viral rebound suggests a greatly reduced pool of target cells at the time of BAM treatment, which was a median of 4.5 days post-symptom onset for the seven individuals analyzed. Thus, a mechanism for a supply of target cells is needed to support the rebound (See S1B and S2, Supplementary Material). We investigate two non-mutually exclusive mechanisms that may contribute to the availability of target cells after peak viral load, i.e., proliferation of target cells (Fang et al., 2020; Liberti et al., 2021; Bridges et al., 2022) and transition of cells from an interferon induced antiviral state back to a susceptible state due to a decline in the level of type I and III interferons (Samuel and Knutson 1982; Ulker and Samuel, 1987; Voigt et al., 2016). Thus, we introduce the modifications needed for each proposed mechanism to explain the rebound of virus.

### Target cell regeneration (the logistic proliferation model)

We assume homeostatic mechanisms induce a logistic growth of target cells that attempts to restore the population size to its baseline level. This leads to the following modification of the target cell dynamics:

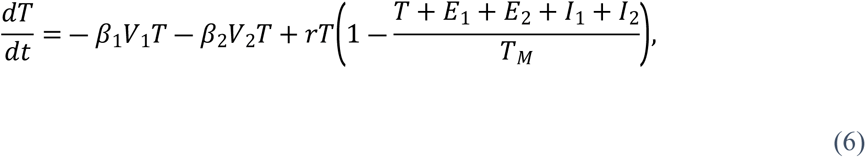

where *r* and *T_M_* are the maximum proliferation rate (or regeneration rate) and carrying capacity, respectively. While it may be possible for infected cells to proliferate and generate infected progeny, existing experimental evidence suggests that proliferation of cells infected with SARS-CoV-2 is impaired (Mizutani et al., 2006; Avendaño-Ortiz et al., 2020; Hekman et al., 2020). Hence, we assume proliferation of infected cells is negligible.

### Loss of the antiviral state (the innate immune response model)

The protective effect of the type-I and type III interferons (representative of the innate immune response) is modeled as in previous studies (Saenz et al., 2010; Pawelek et al., 2012; Ke et al., 2021). We introduce a variable *R* that keeps track of cells in an antiviral state and thus refractory to infection. This leads to the following changes to the original model:

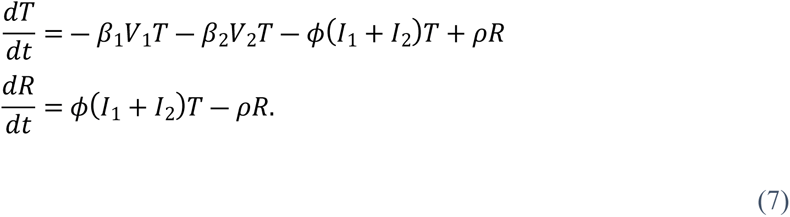

In this formulation, we assume the interferon level is proportional to the number of productively infected cells, so the rate of conversion of target cells to refractory cells is given by the rate constant *ϕ* multiplied by the total number of infected cells, (*I_1_+I_2_*). The refractory cells return to being susceptible to infection at per capita rate *ρ*.

### Population fitting

We used a nonlinear mixed effects modeling approach (software Monolix 2021, Lixoft, SA, Antony, France) to fit each model to the viral load and mutant frequency data for all individuals simultaneously. Specifically, besides fitting the total viral load, we fit 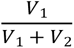 to the frequency of the BAM sensitive viral population. We do not fit 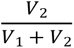 to the frequency of the BAM resistant viral population because it is simply 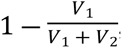, so the resistance data is collinear with the sensitive data. Comparison of the models was done using the corrected Bayesian Information Criterion (BICc) (Burnham and Anderson, 1998, 2004) as reported by Monolix.

Since we do not know the time of infection and estimating the time of infection without data during the early phase of infection is problematic due to model identifiability issues (Ciupe & Tuncer, 2022; Zitzmann et al., 2023), we fixed the duration from infection to symptom onset to 5 days as in previous studies (Gonçalves et al., 2020; Neant et al., 2021; Lingas et al. 2022). This assumption is based on estimates that the duration from exposure to symptom onset is about 5 days (Lauer et al., 2020; He at al., 2020; Linton et al., 2020; Killingley et al., 2022; CDC, 2022). Furthermore, fixing the duration from infection to symptom onset allows us to focus on our objective, which is to access potential mechanisms that contribute to transient viral rebound of resistant virus.

For all models, we fixed the initial number of target cells *T*(*0*) to 8 × 10^7^ cells (Ke et al., 2021). This is an estimate based on the observation that the number of epithelial cells that expresses the ACE-2 receptor is roughly 20% of the epithelial cells in the upper respiratory tract (around 4 × 10^8^ cells) (Hou et al., 2020). For the logistic proliferation model, we set *T_M_* = *T*(0). This means in the absence of viral infection, the proliferation rate is negligible, which is consistent with observation that the turnover of epithelial cells is on the order of months; however, the proliferation rate is accelerated following acute infection (Liberti et al., 2021).

As we do not know the number of viruses that initiate infection, we use a method suggested by Smith et al. (2018) in which we assume the initiating virus is either cleared or rapidly infects cells. Thus, we initiate infection by setting *E*_1_(0) = 1 cell. Also, besides resistance arising by viral mutation there can be baseline resistance. Hence, we fit *E*_2_(0) and keep track of the first time, *t*\*\*, the model predicts *E*_2_(*t*\*\*) ≥ 1, where *t*\*\* marks a physiologically relevant threshold for the emergence of resistance virus that can be compared across individuals.

We fixed the association constant for BAM binding to the SARS-CoV-2 spike protein *k_on_* to 5.5 × 10^5^ M^−1^s^−1^ and the dissociation constant *k_off_* to 2.5 × 10^−5^ s^−1^ (Jones et al., 2021; Jones personal comm.). This gives a dissociation constant (*K_D_*) of 0.045 nM, which is somewhat less than the reported value of 0.071 nM in the EUA for BAM (FDA, 2021). Due to differences in units between the concentration of BAM and these rate parameters, the appropriate units are obtained for the fitting based on the following conversion:

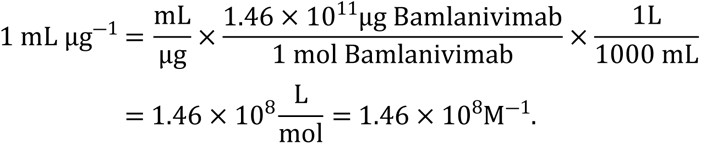

The rate of exit from the eclipse phase, *k,* is fixed to 4 per day and the viral clearance rate, *c*, is fixed to 10 per day (Ke et al., 2021). BAM accelerates viral clearance within three days of infusion (Chew et al., 2022) and thus we expect immune complexes to be cleared faster than free virus. Therefore, we carried out sensitivity tests with *γ* varying between *c* and 5 × *c* based on the observation that neutralizing antibody enhances HIV-1 clearance by ∼3-fold (Igarashi et al., 1999). We found that model fits were relatively insensitive to the value of *γ* in this range and we fixed the clearance rate of the virus-antibody immune complexes to 3 × *c* (see Table S2, Supplementary Material), the value found by Igarashi et al. (1999). The mutation rate *μ* was fixed to 10^−6^ per day (Amicone et al., 2022; Bar-On et al., 2020). Finally, based on the discussion in S2 (Supplementary Material), we also constrained the infection rate constant for the resistant virus *β*_2_ to be less than *β*_1_. A complete summary of parameters is presented in Table S3 (Supplementary Material).

## Results

To demonstrate that target cell replenishment is necessary, we fit a target cell limited model given by Eqs. (1)-(5) to the data. The results of such fit are inconsistent with previous observations about when the viral load peak occurs. The model can recapitulate the second peak, but at the time of treatment, the model predicts that the viral load has yet to reach its normal first peak in all seven participants (Fig. S1, Supplementary Material). This is unlikely since the viral peak is usually around the time of symptom onset and BAM was initiated a median of 4.5 days post symptom onset in these participants (Choudhary., 2022). Essentially, this simple model tries to preserve target cells to allow the resistant virus to grow by preventing the viral load from increasing rapidly after infection. Hence, we proceed with testing models that include target cell replenishment.

### The logistic proliferation model

We first tested the logistic proliferation model, which incorporates proliferation of target cells (see Methods). Fitting the model to data, we show that the model can describe the emergence of resistant virus reasonably well (Fig. 1). Model simulation using the best-fit parameter values recapitulated the observation that the drug sensitive viral population (green in Fig. 1) was the dominant strain in all seven study participants before treatment. After BAM infusion, the sensitive viral population was rapidly replaced by the resistant viral population (red in Fig. 1), which expanded and led to the observed viral rebound.

**Figure 1.**
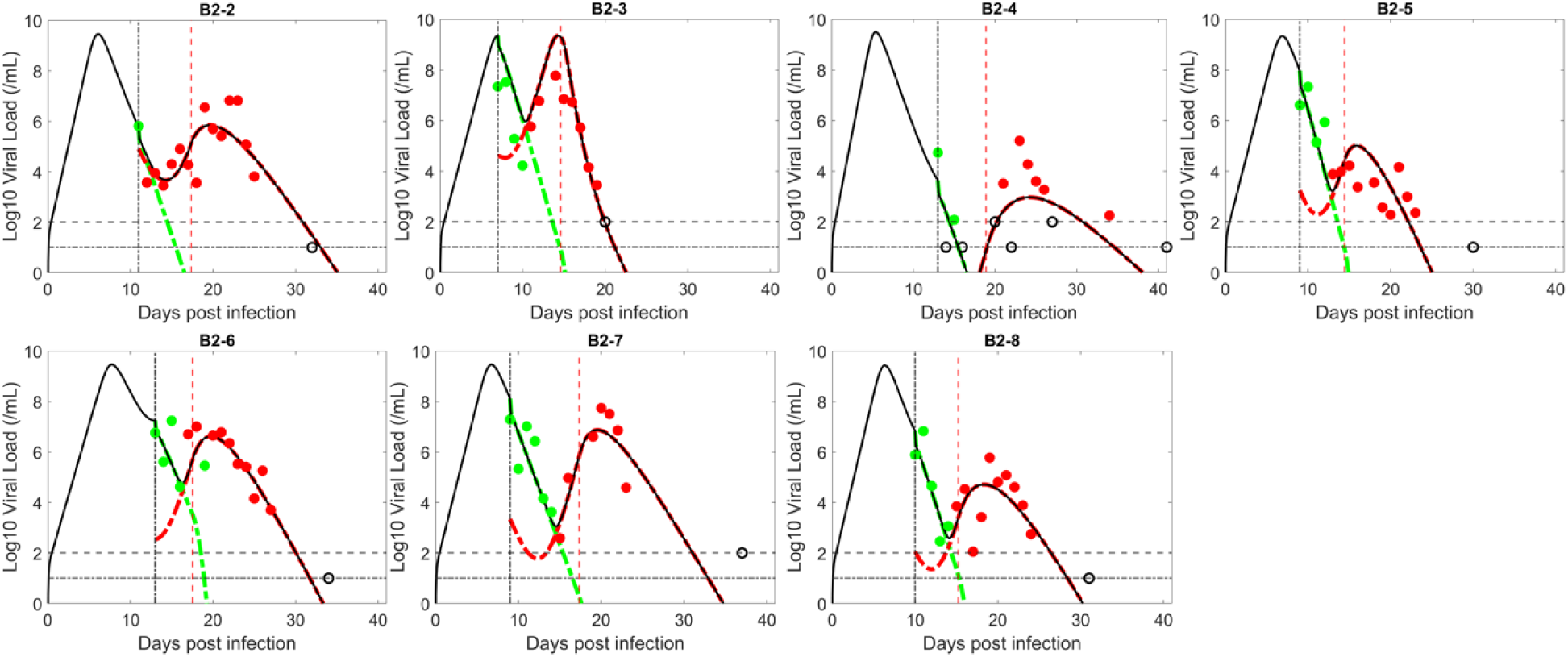
Fit of the logistic proliferation model to the data. The circles represent viral load data which are filled green when the viral population was dominated by BAM sensitive virus and filled red when dominated by resistant virus. The unfilled circles are data below the limit of quantification (2 log_10_ RNA copies/mL) or limit of detection (1.4 log_10_ RNA copies/mL, indicated by horizontal lines). Black curves show the best-fit of the model to the total viral load. When plotting the model fit, *V*_1_ (BAM sensitive) is represented by a dashed–green curve and *V*_2_ (BAM resistant) by a red curve. The vertical black line indicates the time of treatment initiation. The vertical red line indicates the estimated time, *t*\*, when adaptive immunity begins to emerge.

The parameter estimates for the logistic proliferation model are relatively consistent across participants (Table 1). The one exception is the time that the resistant virus emerges *t*\*\*. Participant B2-2 had an estimated resistance emergence time that was much earlier (0.84 days) than the other six people (range 3.3 to 4.8 days). This, however, is consistent with experimental data, which showed B2-2 had baseline (at the time of BAM infusion) resistant mutations E484K and E484Q. For this individual, the anterior nasal swab data (which is used for the fitting), showed baseline mutation frequencies of less than 10%. However, in nasopharyngeal swabs, which were taken much less frequently and hence not used for model fitting, the frequency of the BAM resistance mutation E484K was over 60% at baseline (Choudhary et al., 2022). No baseline mutation was detected in the other six participants in either anterior nasal or nasopharyngeal measurements, which is consistent with the late predicted emergence time of the resistant viral population.

**Table 1.**
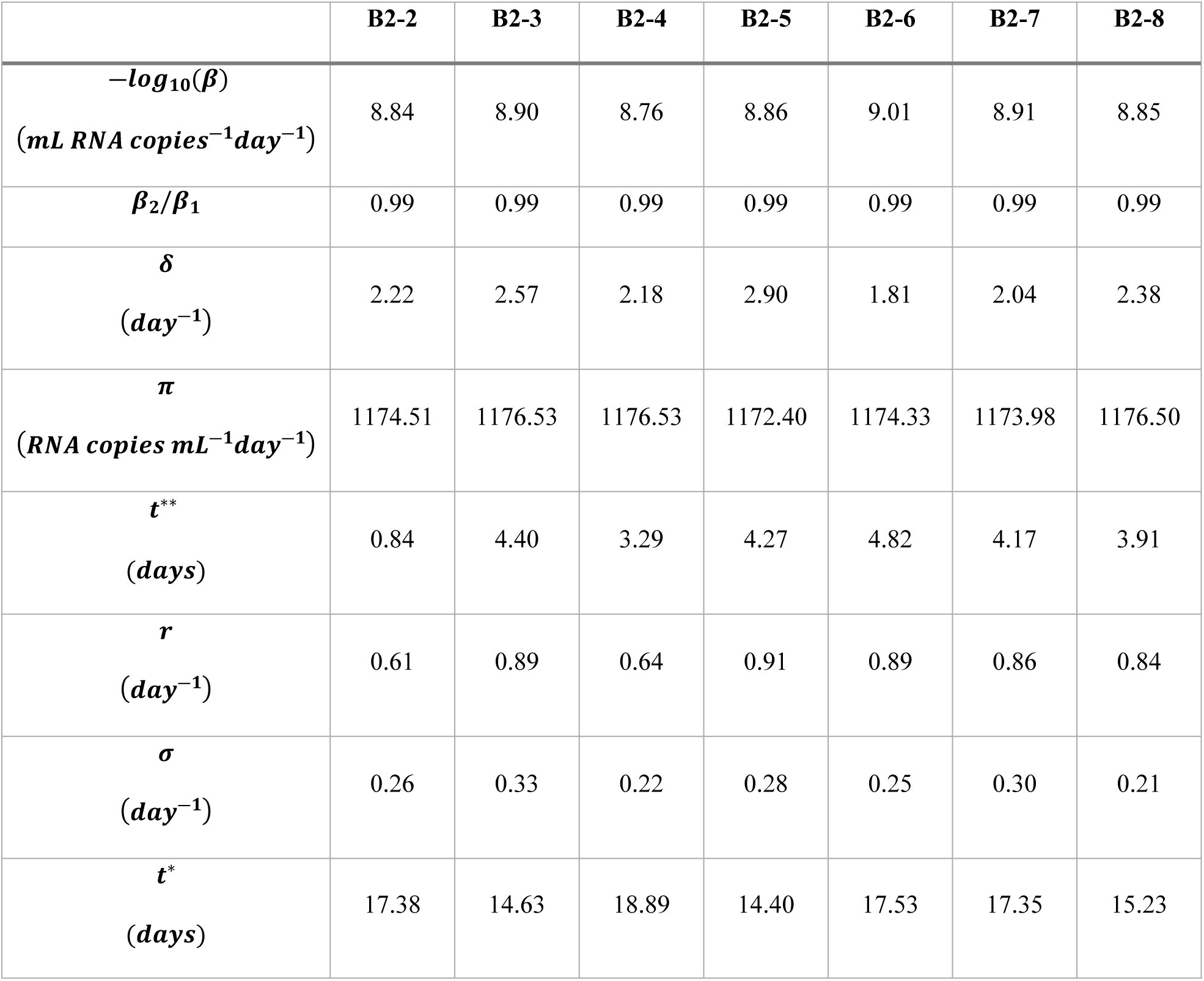
Best fit parameters for the logistic proliferation model.

|  | B2-2 | B2-3 | B2-4 | B2-5 | B2-6 | B2-7 | B2-8 |
| --- | --- | --- | --- | --- | --- | --- | --- |
| $-\log_{10}(\beta)$<br>(*mL RNA copies*<sup>-1</sup>*day*<sup>-1</sup>) | 8.84 | 8.90 | 8.76 | 8.86 | 9.01 | 8.91 | 8.85 |
| $\beta_2/\beta_1$ | 0.99 | 0.99 | 0.99 | 0.99 | 0.99 | 0.99 | 0.99 |
| $\delta$<br>(*day*<sup>-1</sup>) | 2.22 | 2.57 | 2.18 | 2.90 | 1.81 | 2.04 | 2.38 |
| $\pi$<br>(*RNA copies mL*<sup>-1</sup>*day*<sup>-1</sup>) | 1174.51 | 1176.53 | 1176.53 | 1172.40 | 1174.33 | 1173.98 | 1176.50 |
| $t^{**}$<br>(*days*) | 0.84 | 4.40 | 3.29 | 4.27 | 4.82 | 4.17 | 3.91 |
| $r$<br>(*day*<sup>-1</sup>) | 0.61 | 0.89 | 0.64 | 0.91 | 0.89 | 0.86 | 0.84 |
| $\sigma$<br>(*day*<sup>-1</sup>) | 0.26 | 0.33 | 0.22 | 0.28 | 0.25 | 0.30 | 0.21 |
| $t^*$<br>(*days*) | 17.38 | 14.63 | 18.89 | 14.40 | 17.53 | 17.35 | 15.23 |

### The innate immune response model

Next, we tested the ability of the innate immune response model in which target cells arise from the loss of the antiviral state to fit the observed data (see Methods). The model assumes type-I and type-III interferons, which are proportional to the number of infected cells, put target cells into an antiviral state that is refractory to viral infection (Samuel, 2001; Garcia-Sastre and Christine, 2006; Voigt et al., 2016). The antiviral state is not permanent, and the model assumes refractory cells return to being susceptible to infection at a constant rate. As the viral infection progresses, more cells become infected, more interferon is produced, and a larger number of target cells become refractory to infection. However, after the viral load reaches its peak, both the viral load and the number of infected cells decline. As the number of infected cells and interferon levels decline, fewer target cells become protected, while cells in the refractory state lose protection and return to being susceptible to infection. The refractory cells returning to target cells can provide resistant virus with the necessary resource to rebound.

Similar to the logistic proliferation model, the innate immune response model can also capture the emergence of resistant virus and the observed transient viral rebound (Fig. 2). The model also describes well the quantitative dynamics of both the BAM sensitive and resistant viral populations and the observation that prior to treatment, the sensitive virus is dominant, while, after BAM infusion, the resistant viral population rapidly replaces the sensitive one, before finally being cleared by adaptive immunity (Fig. 2).

**Figure 2.**
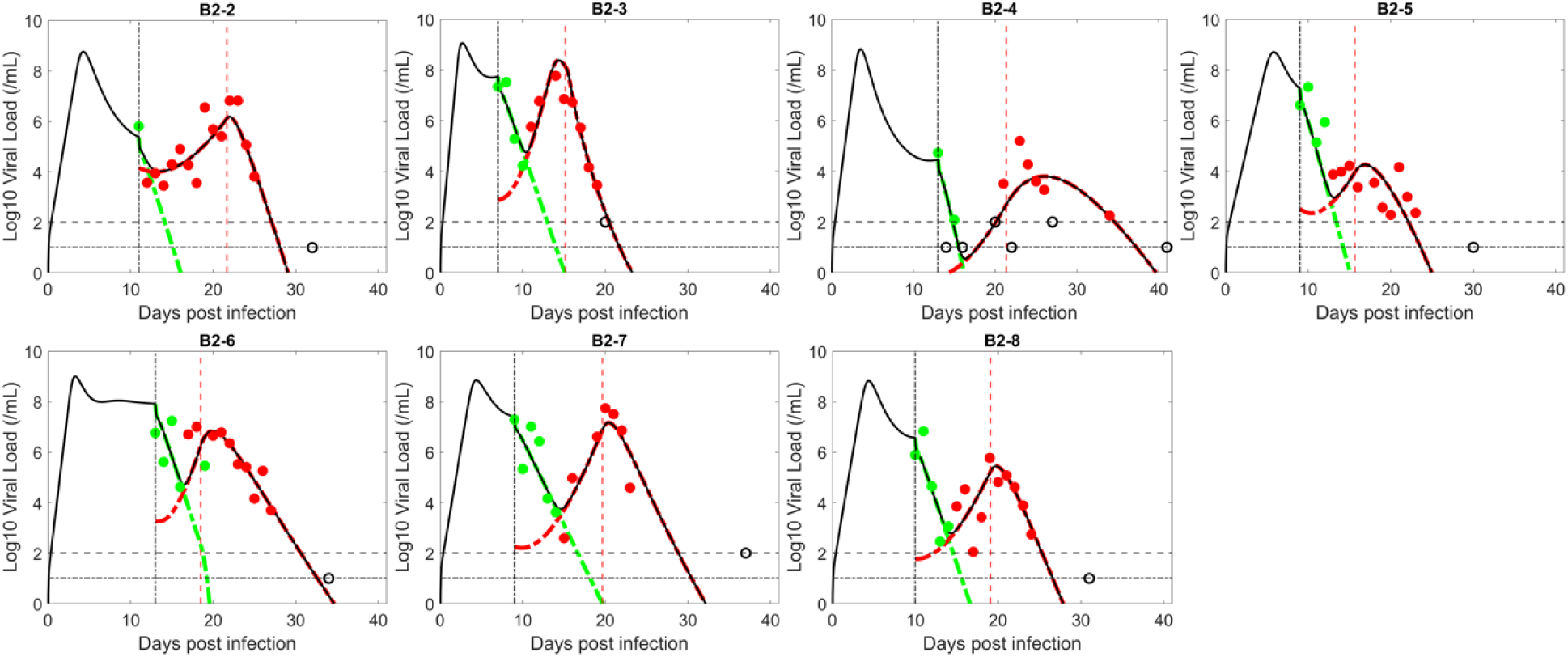
Fit of the innate immune response model to the data. The circles represent viral load data which are filled green when the viral population was dominated by BAM sensitive virus and filled red when dominated by resistant virus. The unfilled circles are data below the limit of quantification or limit of detection (indicated by horizontal lines). Black curves show the best-fit of the model to the total viral load. When plotting the model fit, *V*_1_ (BAM sensitive) is represented by a dashed–green curve and *V*_2_ (BAM resistant) by a red curve. The vertical black line indicates the time of treatment initiation. The vertical red line indicates the estimated time, *t*\*, when adaptive immunity begins to emerge.

The estimated parameters in the innate immune response model are consistent across patients (Table 2). Interestingly, the estimated viral dynamics parameters that are common to both the logistic proliferation and innate response models are very similar (e.g., *β, π* and *δ*). Furthermore, the best estimates of the ratio *β*_2_*β*_1_, in both models, are close to the upper bound of 1, which suggests BAM resistance mutations, such as E484K, have little fitness cost. This is perhaps due to their slightly higher binding affinity to ACE-2 (Barton et al., 2021). The estimated emergence times of the resistant virus for the innate immune response model follow a similar trend to that of the logistic proliferation model, which reflects the observation of a baseline mutation in B2-2 but not the other participants. The estimated values for ρ are consistent with the expected duration of protection of refractory cells (Voigt et al., 2016; Patil et al., 2015; Samuel et al., 1982).

**Table 2.** Best fit parameters for the innate immune response model.

|  | B2-2 | B2-3 | B2-4 | B2-5 | B2-6 | B2-7 | B2-8 |
| --- | --- | --- | --- | --- | --- | --- | --- |
| $-\log_{10}(\beta)$<br>(*mL RNA copies*<sup>-1</sup>*day*<sup>-1</sup>) | 8.61 | 8.20 | 8.40 | 8.79 | 8.39 | 8.67 | 8.63 |
| $\beta_2/\beta_1$ | 0.98 | 0.98 | 0.99 | 0.99 | 0.99 | 0.86 | 0.98 |
| $\delta$<br>(*day*<sup>-1</sup>) | 2.38 | 2.19 | 3.59 | 2.75 | 2.32 | 1.55 | 2.26 |
| $\pi$<br>(*copies mL*<sup>-1</sup>*day*<sup>-1</sup>) | 1097.54 | 1102.81 | 1102.53 | 1102.11 | 1108.76 | 1100.88 | 1102.17 |
| $\phi$<br>(*cells*<sup>-1</sup>*day*<sup>-1</sup>) | 8.1e-07 | 9.5e-07 | 8.1e-07 | 4.8e-07 | 7.5e-07 | 7.8e-06 | 6.9e-07 |
| $\rho$<br>(*day*<sup>-1</sup>) | 0.021 | 0.069 | 0.021 | 0.1 | 0.15 | 0.053 | 0.04 |
| $t^{**}$<br>(*days*) | 0.77 | 1.61 | 2.24 | 3.87 | 2.04 | 2.96 | 2.85 |
| $\sigma$<br>(*day*<sup>-1</sup>) | 0.32 | 0.65 | 0.038 | 0.12 | 0.27 | 0.41 | 0.31 |
| $t^*$<br>(*day*) | 21.65 | 15.16 | 21.37 | 15.66 | 18.48 | 19.65 | 19.1 |

We compare the data support for each model using the negative log-likelihood (-2LL), the AIC and BICc for both models (Table 3). The innate immune response model fits the data slightly better than the logistic proliferation model (lower fitting error); however, the logistic proliferation model is slightly preferred based on BICc, because the innate immunity model has one more model parameter, which is two more estimated parameters since population fitting estimates a mean and a standard deviation for the parameter.

**Table 3.** Comparison of fitting between the logistic proliferation and innate immune response models. Bolded text shows the smaller value.

|  | -2LL | BICc |
| --- | --- | --- |
| Logistic proliferation model | 191.59 | **260.63** |
| Innate immune response model | **190.68** | 267.02 |

### Adaptive immunity is crucial in viral clearance

In a target cell limited model (see S2 Supplementary Material), the constant depletion of target cells ultimately leads to the decline in viral load; however, both the logistic proliferation model and the innate immune response model contain mechanisms that can replenish the pool of target cells. In a logistic proliferation model without adaptive immunity, the production of target cells makes it possible for virus to oscillate or be sustained indefinitely (see text S4, Supplementary Material). In the innate immune response model without adaptive immunity, the dynamics of target cells going in and out of the refractory state can also give rise to oscillations and the possibility of a slowly decreasing viral load (see Fig. S2 and text S4, Supplementary Material). Thus, for these models to generate dynamics consistent with the observed data with rapid ultimate virus clearance, adaptive immunity must be included in both models.

### Variations in individual characteristics may explain why only some develop resistance with high transient viral rebound

A previous study using a similar innate immune response model for SARS-CoV-2 infection but fit to a different set of data from untreated, unvaccinated individuals estimated *ϕ* = 3.4 × 10^−6^ mL/day and *ρ* = 0.004/day (mean of the estimates for all individuals studied) (Ke et al., 2021). Compared to these estimates, our estimate for *ϕ* is slightly smaller, but within one order of magnitude; however, our estimate for *ρ* is at least one order of magnitude larger. This difference in the rate that refractory cells lose protection, *ρ,* along with the ability of BAM to neutralize wildtype virus may explain why the viral trajectories for these seven study participants exhibit high transient viral rebound of resistant virus (Choudhary et al., 2022).

To study this further, we used the best fit parameters for B2-8 and simulated the effect of decreasing the supply rate of target cells, *r* (the logistic proliferation model) or the rate cells lose their antiviral state *ρ* (the innate immune response model). We found that decreasing the maximum target cell proliferation rate *r* in the logistic proliferation model caused the magnitude of the transient viral rebound to diminish (Fig. 3, left). Reducing the rate that refractory cells return to the target cell pool also has a similar effect (Fig. 3, right). However, there is a distinction in the qualitative effect of the two mechanisms. The proliferation term creates newly available target cells to compensate for the number of cells lost, while the refractory term only stores target cells in an antiviral state temporarily and does not create new target cells. These results suggest that how quickly target cells become available during infection may be a key factor determining the magnitude of the transient viral rebound associated with the emergence of resistant virus. Since these parameters vary among individuals, it may explain why only some individuals exhibit high resistant viral rebound following treatment with BAM. Additionally, differences in mutations in spike, differences in BAM binding kinetics, differences in BAM concentration in different treated individuals, differences in the time BAM was given relative to time of infection, differences in viral kinetics, and in particular higher viral load at the time of BAM treatment in participants with treatment-emergent resistance than those who did not develop resistance (Choudhary et al., 2022), etc. may further contribute to the observation that only a small fraction of treated individuals develop resistance and transient viral rebound.

**Figure 3.**
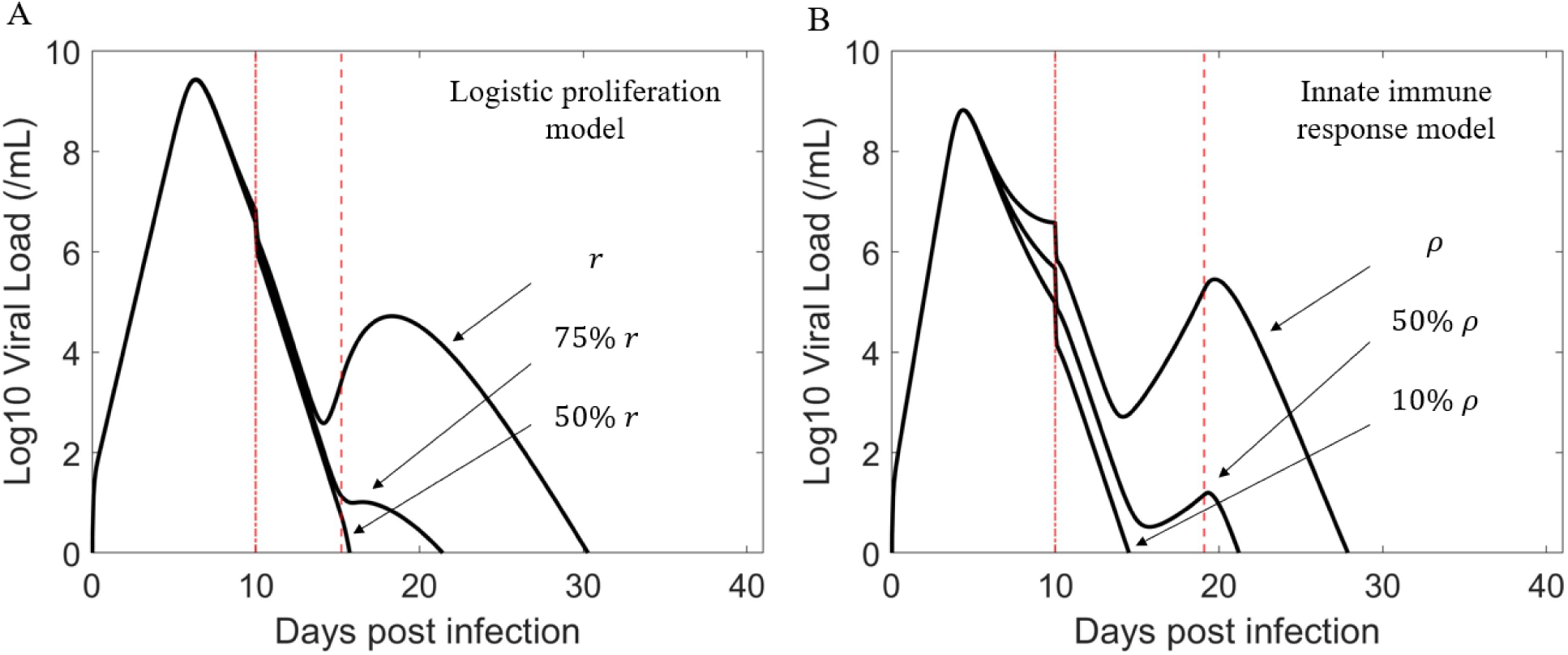
The rate of target cell replenishment has a crucial role in driving the amplitude of the viral rebound. Baseline parameters for the simulation are taken from the best fit parameters of B2-8. The first vertical red (dashed dot) line indicates the time of treatment initiation. The second vertical red (dashed) line indicates when adaptive immunity begins to emerge. (A) Viral rebound is more likely to be observable (e.g., sufficiently high VL) with increasing intrinsic growth rate *r*. (B) Viral rebound is more likely to be observable with increasing rate of refractory cells returning to cells susceptible to infection *ρ*.

## Discussion

The emergence of BAM resistant viral mutants associated with transient viral rebounds of 3-4 viral logs has been observed in some individuals treated with BAM (Choudhary et al., 2022; Jensen et al., 2021; Peiffer-Smadja et al., 2021). While the emergence of resistant viral mutants can be attributed to the selective pressure of BAM, the 3-4 log increase in viral load is also likely dependent upon having an additional supply of target cells. Viral dynamic models without target cell replenishment are suitable to describe acute infection (Gonçalves et al., 2021b; Kim et al., 2021). However, we found that simple target cell limited models were not consistent with the data (Fig. S1). To observe the large increase in viral load with the emergence of resistance, such models predict treatment takes place early during infection (before peak in viral load), which was not the case here. To reconcile these contradicting observations and to better understand the factors leading to the emergence of resistant virus with high transient viral rebound, we developed two models with different underlying mechanisms that allow for target cell replenishment during the infection.

In an acute infection, target cells production is often assumed to be negligible due to the short duration of the infection—continued viral infection is then dependent upon the availability of target cells (Perelson and Ke, 2021). However, emerging resistant virus with viral rebound extends the duration of infection, which may justify the inclusion of target cell proliferation in the model. Previous SARS-CoV-2 models examined the effect of proliferation of target cells, but not in light of the emergence of resistance and transient viral rebound (Asher et al., 2022; Fatehi et al., 2021; Hattaf et al., 2020; Chatterjee et al., 2020; Du and Yuan, 2020; Li et al., 2020). We showed that by incorporating a logistic proliferation term into a two-viral population viral dynamic model with adaptive immunity, the model can describe the emergence of resistant virus, which leads to the high transient viral rebound following treatment with BAM (Fig. 1). This result echoes a previous observation that in mice with influenza, stimulation of alveolar type II cells prior to infection can increase the number of alveolar epithelial cells expressing ACE-2 receptors and enhance infection (Nikolaidis et al., 2017).

Upon viral infection, infected cells as well as plasmacytoid dendritic cells (not modeled), release type-I and type-III interferons, which put neighboring target cells into an antiviral state, protecting them from viral infection (Talemi and Höfer, 2018; Voigt et al., 2016; Garcia-Sastre and Christine, 2006; Samuel, 2001). Thus, a strong interferon response correlates with strong suppression of viral replication (Jamilloux et al., 2020). Previous modeling efforts have captured these effects in a different context (Ke et al., 2021; Pawelek et al., 2012; Padmanabhan et al., 2022). However, over time, refractory cells likely lose their antiviral state (Voigt et al., 2016). Thus, if there are sufficient numbers of cells in the refractory state and these return to being susceptible to infection, then the returning cells would contribute to a late increase in the pool of susceptible cells. In this case, if individuals were treated with an mAb that effectively neutralizes the sensitive virus, then it would create an opportunity for the resistant virus to infect the increasing number of target cells, allowing the vigorous growth of resistant virus associated with an observable transient viral rebound.

SARS-CoV-2 may be able to antagonize the initial innate immune response by inducing lower production of type I and III interferons as compared to other respiratory viruses, which has been linked to the severity of SARS-CoV-2 disease (Blanco-Melo et al., 2020). However, the importance of type-I interferons in SARS-CoV-2 infection has been highlighted by the finding that at least 10% of people with life-threatening COVID-19 pneumonia had neutralizing auto-antibodies against type I interferon (Bastard et al., 2020). There is a sharp increase in the presence of such antibodies with age in the elderly and people with these auto-antibodies account for at least 18% of all COVID-19 fatalities (Bastard et al., 2021), again highlighting the protective role of an interferon response.

Two SARS-CoV-2 studies using a murine model (Israelow et al., 2020, 2021) suggested that innate immunity may only have a minimal role in viral clearance, while adaptive immune significantly contributes to the ultimate clearance of acute infection. A recent clinical study also echoed this result by examining the role of CD4+ T-cells in shaping the antiviral activity of the immune response (Dong et al., 2022). We have also shown that the inclusion of adaptive immunity in our models is crucial for viral clearance (Fig. S2).

The supply rate of target cells should have a strong impact on the magnitude of the transient viral rebound of resistant virus, or whether the emergence of resistant virus is observable clinically. We examined this effect by varying the target cell supply rate in both models. Figure 3 confirmed our speculation. This suggests that how quickly target cells become available during infection may be a key factor along with BAM driving the high transient viral rebound of resistant virus. Because the target cell supply rate most likely varies among individuals, this may help explain why only some individuals treated with BAM develop observable resistant viral rebound. The innate immune response model predicts that the rapid reduction in infected cells due to BAM led to a reduction in the production of type-I and type-III interferons. With less interferon fewer target cells are converted into the refractory state. In this scenario, if the rate that refractory cells return to susceptible cells is sufficiently fast, it would lead to a sizable increase in the number of target cells that allow for the resistant virus to emerge to an observable population size. While BAM selects for resistant viral population, target cell replenishment drives the amplitude of the transient viral rebound. See Figure 4 for a summary description of our results

**Figure 4.**
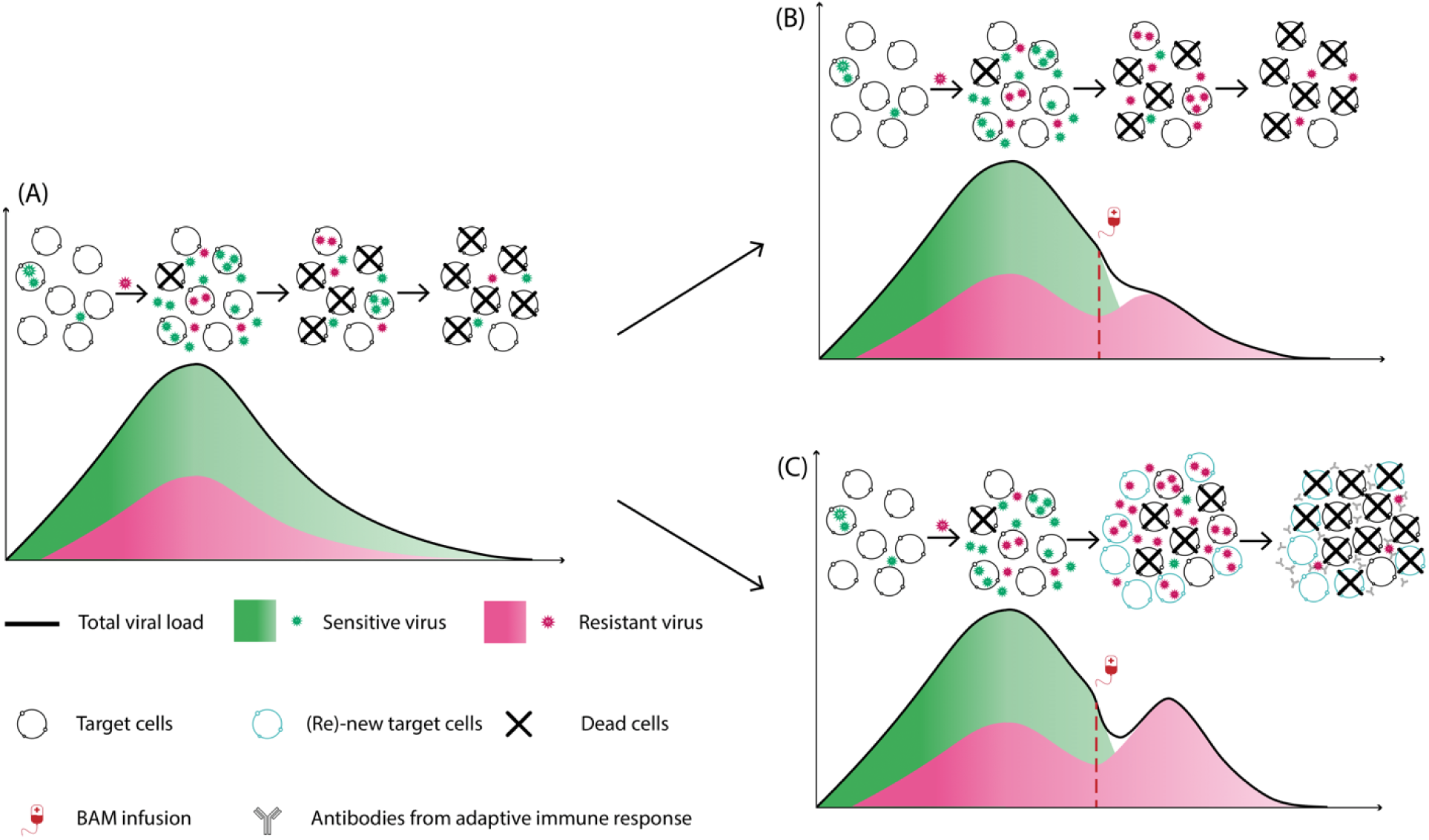
Target cell replenishment can explain the emergence of resistant virus associated with high transient viral rebound. (A) Natural course of acute infection as described by a standard viral dynamic model. (B) Without target cell replenishment, the resistant virus becomes the dominant population but does not lead to observable transient viral rebound. (C) Target cell replenishment – either via new production or target cells return from the refractory state - can drive the resistant viral population to an observable transient viral rebound.

In addition to the above two mechanisms, we also examined the possibility of antibody-dependent enhancement (ADE) extending the range of cells that are targets of infection (not shown). Recent studies provide evidence that SARS-CoV-2 -antibody immune complexes can infect and replicate in macrophages via interactions with FcγRIIA and FcγRIIIA receptors, or even via the complement component C1q receptor (Junqueira et al., 2020; Maemura et al., 2021; Okuya et al., 2022; Zhou et al., 2021; Wu et al., 2020). Based on these findings, we can hypothesize that if the virus-antibody immune complexes are not cleared at a sufficiently fast rate, their ability to infect additional cell types via ADE may allow the resistant strain to take over, leading to the observed transient viral rebound. However, data showing that ADE is important in vivo is lacking and one recent study showed that although an infection-enhancing effect of human neutralizing antibodies was observed in vitro, the same neutralizing antibodies exhibited a protective effect in vivo, when infused into mice or macaques (Li et al., 2021). Another recent analysis using both in vitro and in vivo experiments involving BAM also found no support for it generating ADE of SARS-CoV-2 infection (Cross et al., 2022).

In this study, we tested the hypothesis that the generation of drug resistant variants coupled with an additional supply of target cells can lead to the observed transient viral rebound in some persons with acute SARS-CoV-2 infection following treatment with BAM. Two models were formulated with different mechanisms that allow for the replenishment of target cells during the infection. By fitting the models to patient data, we showed that both models can explain the emergence of resistant virus associated with large transient viral rebounds of 3-4 logs. Comparison between the two models showed that the logistic proliferation model was slightly preferred based on BICc due to having fewer parameters to fit. Nevertheless, both mechanisms may be operational. We also found that variations in the target cell supply rate parameters have a strong impact on the magnitude of the viral rebound, which may explain why only some individuals develop observable transient resistant viral rebound. Finally, we highlighted the role of adaptive immunity in viral clearance in our models.

## Acknowledgements

This work was performed under the auspices of the US Dept. of Energy under contract 89233218CNA000001 and supported by NIH grant U54-HL143541 (RK), UM1AI068634, UM1AI068636 and UM1AI106701, and Los Alamos National Laboratory LDRD 20200743ER (RMR), 20200695ER (ASP), 20210730ER (RMR) and 20220791PRD2 (TP). The authors thank the study participants, site staff, site investigators, and the entire ACTIV-2/A5401 study team.

## Author contributions

**Conceptualization:** Tin Phan, Ruian Ke, Ruy M. Ribeiro, Alan S. Perelson.

**Data curation:** Kara W. Chew, Davey M. Smith, Michael D. Hughes, Manish C. Choudhary, Rinki Deo, Eric S. Daar, David A. Wohl, Joseph J. Eron, Judith S. Currier, Jonathan Z. Li.

**Formal analysis:** Tin Phan.

**Funding acquisition:** Ruian Ke, Ruy M. Ribeiro, Alan S. Perelson.

**Investigation:** Tin Phan, Carolin Zitzmann, Kara W. Chew, Davey M. Smith, Michael D. Hughes, Manish C. Choudhary, Rinki Deo.

**Methodology:** Tin Phan, Ruian Ke, Ruy M. Ribeiro, Alan S. Perelson.

**Project administration:** Ruian Ke, Ruy M. Ribeiro, Alan S. Perelson.

**Resources:** Kara W. Chew, Davey M. Smith, Michael D. Hughes, Manish C. Choudhary, Rinki Deo, Eric S. Daar, David A. Wohl, Joseph J. Eron, Judith S. Currier, Jonathan Z. Li, Ruian Ke, Ruy M. Ribeiro, Alan S. Perelson.

**Software:** Tin Phan, Carolin Zitzmann.

**Supervision:** Ruian Ke, Ruy M. Ribeiro, Alan S. Perelson.

**Validation:** Tin Phan, Ruian Ke, Ruy M. Ribeiro, Alan S. Perelson.

**Writing – original draft:** Tin Phan, Carolin Zitzmann, Ruian Ke, Ruy M. Ribeiro, Alan S. Perelson.

## Writing – review & editing

Tin Phan, Carolin Zitzmann, Kara W. Chew, Jonathan Z. Li, Ruy M. Ribeiro, Ruian Ke, Alan S. Perelson.

## Data Availability

No original data was generated for the paper. The viral loads and the primary resistance mutation frequencies from anterior nasal swabs for the study subjects are all available in the Extended Data section of Choudhary et al. (2022).

